# An LTR retrotransposon in the promoter of a *PsMYB10.2* gene associated with the regulation of fruit flesh color in Japanese plum

**DOI:** 10.1101/2022.01.26.477575

**Authors:** Arnau Fiol, Sergio García, Christian Dujak, Igor Pacheco, Rodrigo Infante, Maria José Aranzana

## Abstract

Japanese plums exhibit wide diversity of fruit coloration. The red to black hues are caused by the accumulation of anthocyanins, while their absence results in yellow, orange or green fruits. In *Prunus, MYB10* genes are determinants for anthocyanin accumulation. In peach, QTLs for red plant organ traits map in an LG3 region with three *MYB10* copies (*PpMYB10*.*1, PpMYB10*.*2* and *PpMYB10*.*3*). In Japanese plum the gene copy number in this region differs with respect to peach, with at least three copies of *PsMYB10*.*1*. Polymorphisms in one of these copies correlate with fruit skin color. The objective of this study was to determine a possible role of LG3-*PsMYB10* genes in the natural variability of the flesh color trait and to develop a molecular marker for marker-assisted selection (MAS). We explored LG3-*PsMYB10* variability, including the analysis of long-range sequences obtained in previous studies through CRISPR-Cas9 enrichment sequencing. We found that the *PsMYB10*.*2* gene was only expressed in red flesh fruits. Its role in promoting anthocyanin biosynthesis was validated by transient overexpression in Japanese plum fruits. The analysis of long-range sequences identified an LTR retrotransposon in the promoter of the expressed *PsMYB10*.*2* gene that explained the trait in 93.1% of the 145 individuals analyzed. We hypothesize that the LTR retrotransposon may promote the *PsMYB10*.*2* expression and activate the anthocyanin biosynthesis pathway. We provide a molecular marker for the red flesh trait which, together with that for skin color, will serve for the early selection of fruit color in breeding programs.

## INTRODUCTION

Anthocyanins are plant pigments synthesized from the phenylpropanoid pathway and they confer red, purple or almost black coloration based on their structural conformation and accumulation levels^1^. These flavonoids protect plants against biotic and abiotic stress, and also enhance animal-mediated flower pollination and seed dispersion^2^. Some edible fruits are a good source of anthocyanins and their color is considered a fruit quality parameter, not only because of the visual attractiveness to the consumer, but also because anthocyanin ingestion has been correlated to several beneficial effects on human health^3^. The Rosaceae family includes species with edible fruits such as apples, pears, strawberries, peaches, cherries, apricots and plums, among others. Some of them have tremendous intraspecific fruit color variability, with anthocyanins conferring color to the whole fruit or single tissues, or not contributing to the final coloration at all.

The R2R3-MYB genes are among the largest transcription factor gene families in plants. They mostly regulate plant-specific processes, including anthocyanin biosynthesis by acting as transcriptional activators as well as repressors^4,5^. In rosaceous species, *R2R3-MYB10* genes have been found to regulate anthocyanin levels in the fruit^6,7^. R2R3-MYB proteins regulate anthocyanin biosynthesis through interaction with a basic helix-loop-helix (bHLH) transcription factor and a WD-repeat protein, which results in the MBW complex that can bind to the promoters of several anthocyanin biosynthesis structural genes and enhance their expression^8-10^. Other R2R3-MYBs downregulate anthocyanin biosynthesis by diverging to other branches of the phenylpropanoid pathway or by repressing structural genes^11-13^. However, the *MYB10* genes remain the main factors leading to red-to-blue fruit coloration, and DNA polymorphisms with direct impact on its expression or protein function have been identified and correlated with fruit color variation in strawberry^14^, apple^15,16^, pear^17,18^, peach^19,20^, sweet cherry^21,22^ and Japanese plum^23^. Transposons have also been reported to play a role in this variability^14,24^. Anthocyanin levels can also be modulated by ethylene production^25^ and environmental factors such as light and temperature^26-28^, adding an extra layer of complexity to their regulation.

Anthocyanins are synthetized in many plant tissues, constitutively or under certain conditions. In the *Prunus* genome, several QTLs for fruit color have been mapped in its eight linkage groups (LG), some of them in agreement with the position of a cluster of MYB10 genes in LG3^29-31^. In peach, this cluster contains three MYB10 genes (*MYB10*.*1, MYB10*.*2* and *MYB10*.*3*)^32^, while an increased number of MYBs have been annotated in the syntenic region of the sweet cherry, apricot and Japanese plum genomes^33-35^.

The multiple loci mapped in the *Prunus* genome for anthocyanin accumulation and the complexity of the phenotype highlight the complex regulation of synthesis of this pigment, in which several genes, differing in their spatial and temporal expression patterns, promote or repress the phenylpropanoid pathway. In peach, the recessive blood flesh (*bf*), mapped in LG4, promotes the accumulation of anthocyanins in the flesh after the pit hardening stage, i.e. long before fruit maturity^36^, while the dominant blood flesh (*DBF*), mapped in LG5, controls the blood-flesh trait at the final stages of fruit maturation^37^. The flesh color around the stone (*Cs*) mapped in LG3^38^ explains a localized pigment accumulation, unlike the highlighter (*h*) in the same LG that suppresses anthocyanin accumulation in the whole fruit^20,39^. Some of these traits, such as DBF, are regulated through the activation of MYB10 gene expression^40^, while others are determined by polymorphisms in the MYB10 genes themselves^19,20,41^.

Japanese plum (*Prunus salicina* L.) is one of the rosaceous crops with more fruit color variability. Fiol et al.^23^ studied the allelic variability in the LG3-*PsMYB10* region. They found a triplication of the *PsMYB10*.*1* gene in some of the varieties and identified an allele in one of the gene copies highly associated with the skin anthocyanin color. In addition, they characterized the variability into haplotypes, providing an efficient molecular marker for marker-assisted selection (MAS) of skin color in breeding programs. However, they found no allele significantly associated with the red flesh color. To our knowledge, only two studies have found polymorphisms, mapped in LG6 and LG3, associated with the fruit flesh color in Japanese plum^42,43^. The number of individuals used in both studies was reduced, which limits the transfer of the markers to broader and more variable collections and motivates the search for markers more tightly linked to the trait.

The well-known role of *MYB10* genes in rosaceous fruit color variation the, weak association found between LG3 makers and flesh color in Japanese plum and the diverse number of alleles and gene copies identified by Fiol et al.^23^ motivated a deeper exploration of the genetic variability of the LG3-*PsMYB10* region. For that, Fiol et al.^44^ used CRISPR-Cas9 target enrichment sequencing in a pool of Japanese plum varieties, confirming the high level of variability in the MYB10 region between Japanese plum varieties compared to other *Prunus* genomes; and finding that the level of homology between Japanese plum varieties was comparable to that between *Prunus* species and their wild relatives. The CRISPR-Cas9 target sequencing approach yielded an excellent source of polymorphisms (SNPs and SVs) that can be used to find additional variants associated with fruit color, especially in the flesh.

The objective of this study was to explore the LG3-*PsMYB10* variability in the search for polymorphisms associated with the red flesh trait to further design a molecular marker for MAS in Japanese plum. By using the available MYB10 molecular marker provided in Fiol et al.^23^, we found the association of one haplotype (H2) whose *PsMYB10*.*2* gene was expressed only in the mature flesh of red fruits. We validated the role of *PsMYB10*.*2* in the activation of anthocyanin biosynthesis by its transient expression in yellow fruits. After exploring the *de novo* sequences from the CRISPR-Cas9 experiment in Fiol et al.^44^, we identified a copia-like LTR retrotransposon inserted in the promoter of the *PsMYB10*.*2* expressed genes only. The presence/absence of the LTR retrotransposon explained 93.1% of the red flesh color trait acquired during the last stage of ripening, becoming an efficient molecular marker for MAS in Japanese plum breeding programs.

## RESULTS

### The MYB10-H2 haplotype explains most of the flesh color variability

We studied the correlation between the LG3-MYB10 haplotypes described by Fiol et al.^23^ and the fruit flesh color in a sample of 103 Japanese plum selections (Figure 1) (Supplementary Data 1). A χ^2^ test found haplotype H2 highly correlated with red flesh (p-value=8.06×10^−12^) (Supplementary Table 1). Forty-one produced fruits with red flesh, 33 of them (80.5%) with H2 in either homozygosis or heterozygosis. Haplotype H2 was present in seven non-red selections (11.29% in the non-red group), including one that was homozygous (C57, H2/H2). Five of them had an infrequent allele (a467) not observed in the H2 of red flesh selections (see Fiol et al.^23^ for more details), indicating that there may be two different H2 haplotypes.

**Figure 1.**
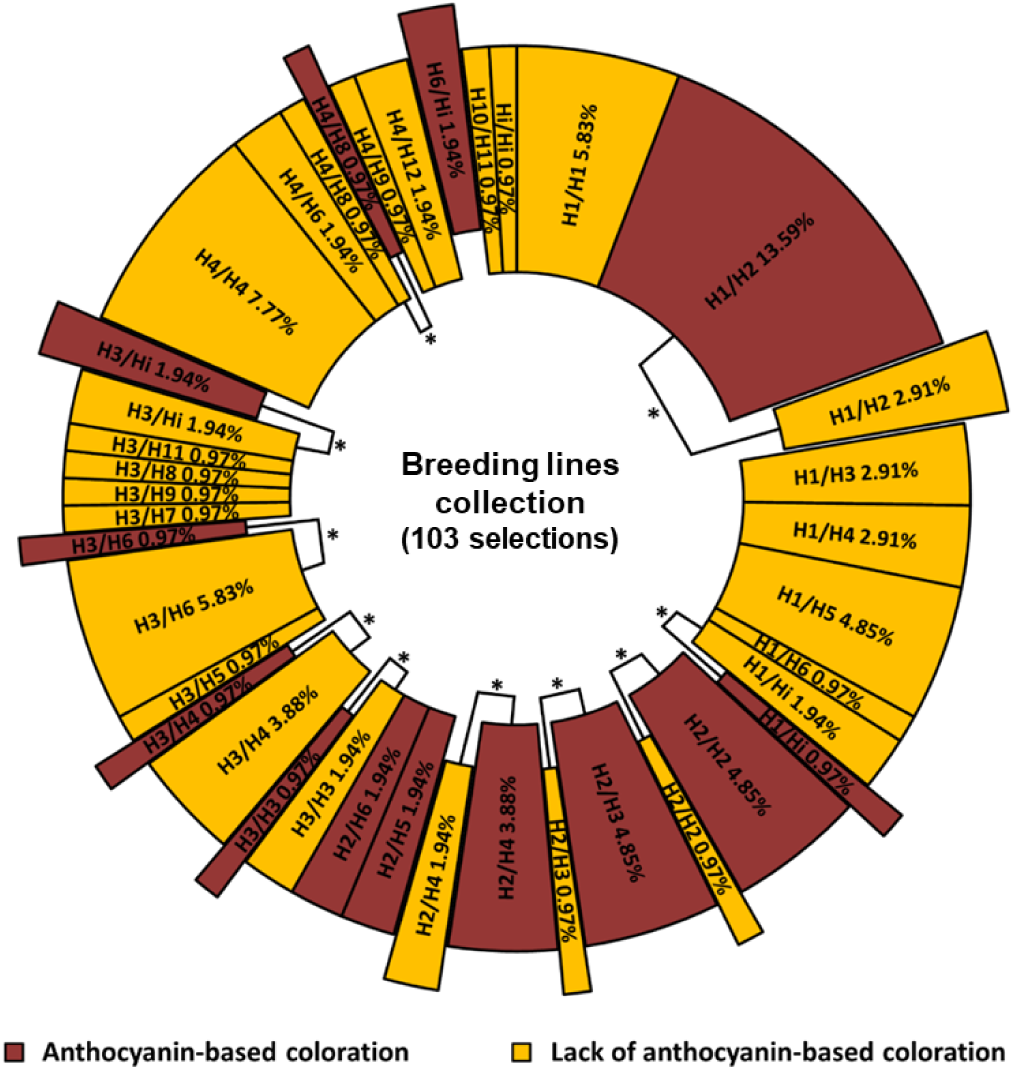
Graphic representation of haplotype combinations and flesh color in 103 Japanese plum selections. Portion size is proportional to the number of individuals. Ten of the haplotype combinations were found in both red and yellow fruits (*). Haplotype H2 was associated with the red flesh color in 84.47% of the tested genotypes; individuals with phenotype not explained by H2 (15.53%) are displaced out of the circle.

### The PsMYB10.2 gene is expressed in the red flesh

The haplotype H2 defined in Fiol et al.^23^ was characterized by three *PsMYB10* alleles: a470 homologous to *PsMYB10*.*1*, a466 homologous to *PsMYB10*.*2*, and a492 homologous to *PsMYB10*.*3*. None of these alleles were exclusive to H2. We tested their expression in the flesh of mature fruits of five varieties with different flesh color and haplotype combinations. Only the *PsMYB10*.*2* allele (a466) was expressed in the flesh, and only in the red-fleshed fruits (Figure 2). The *PsMYB10*.*2* sequence amplified from mature fruit flesh cDNA had an approximate length of 800 bp, while the same primers amplified a fragment of around 2 kb in genomic DNA.

**Figure 2.**
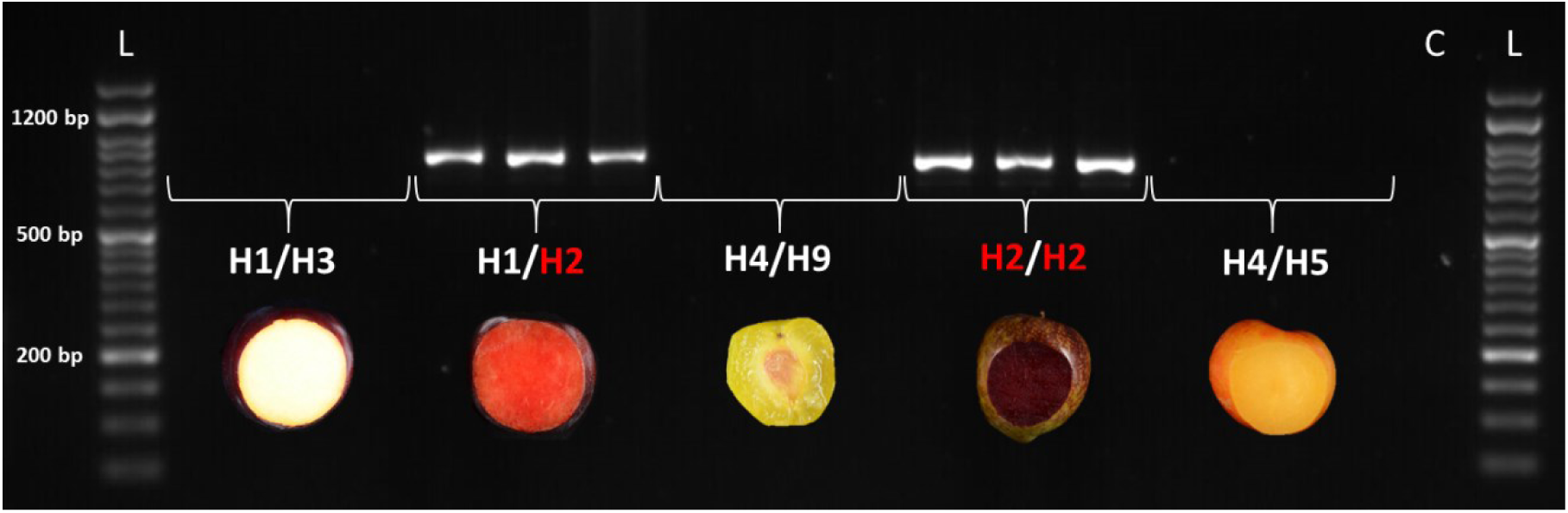
Expression of the *PsMYB10*.*2* gene in the mature flesh of five Japanese plum varieties (three fruits per variety). H1, H2, H3, H4, H5 and H9 are LG3-MYB10 haplotypes defined in Fiol et al.^23^. Well L: 50 bp ready-to-use DNA Ladder (GeneON); well C: no template control reaction.

### The predicted PsMYB10.2 protein sequence has low variability

The RT-PCR bands, as well as the *PsMYB10*.*2* bands from genomic DNA of H1 to H4, were sequenced from C3, C31, C6 and C12, respectively. A complete match of the RT-PCR sequences with the exon sequences of the *PsMYB10*.*2-H2* gene was observed. The alignment of the predicted protein sequences of the *PsMYB10*.*2* alleles in the haplotypes H1 to H4 as well as that predicted from the NCBI *PsMYB10*.*2* GenBank sequence EU155161.1 revealed variation in some aminoacids (Figure 3). While the *PsMYB10*.*2*-H2 and H4 alleles coded for proteins with 243 amino acids; the *PsMYB10*.*2-* H1 and H3 alleles had three additional amino acids in their C-terminal sequence. All predicted proteins had homology higher than 95%, the proteins of H2 and H3 being the closest, with 99.18% homology, while H4 was slightly more distant to all the others, being 95.06% homologous with H1 and 95.88% with H2 and H3 (Supplementary Data 2). The amino acid sequences were also compared with the peach PpMYB10.2 translated protein (*Prupe*.*3G163000*) (Figure 3), where the functionality is significantly reduced due to the simultaneous occurrence of a lysine (K) and an arginine (R) at positions 63 and 90, respectively^45^. In these positions, all the PsMYB10.2 proteins were monomorphic, with an arginine (R) in both positions. Motifs and other key amino acids reported by Stracke et al.^5^, Lin-Wan et al.^7^ and Yang et al.^46^ were identified; none of the amionacid polymorphisms between the PsMYB10.2 proteins occurred at these positions.

**Figure 3.**
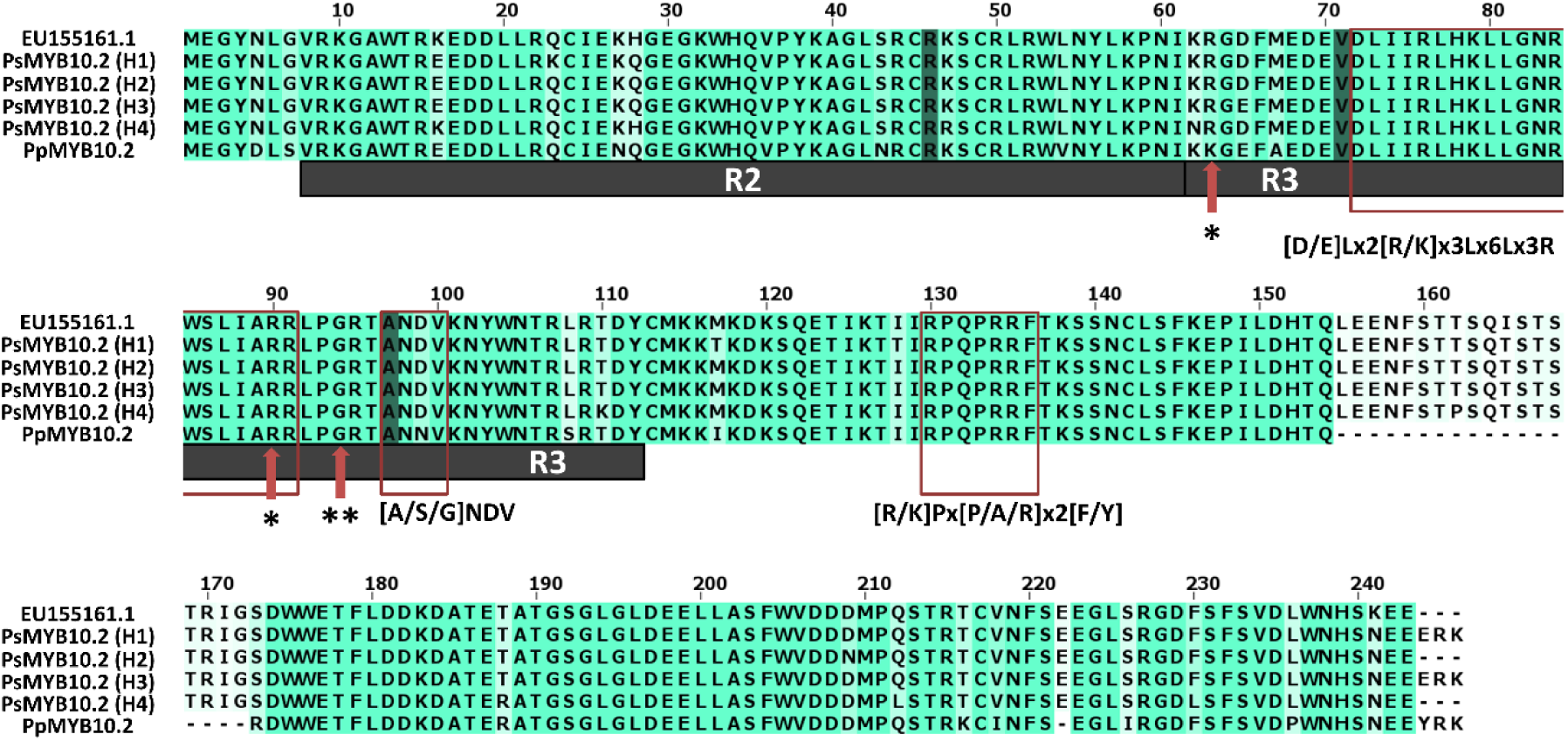
Alignment of the *in silico* proteins of the *PsMYB10*.*2* genes on haplotypes H1 to H4, the Japanese plum *PsMYB10*.*2* from the NCBI database (EU155161.1) and the peach *MYB10*.*2* (*PpMYB10*.*2*) from the Peach Genome v2^47^. The conserved R2 and R3 domains are shown. The motifs from orthologous MYBs (red rectangles) and the conserved residues from anthocyanin-promoting R2R3-MYBs from the Rosaceae family (shadowed residues) were monomorphic in the four haplotypes. Red arrows indicate key function residues reported by (*) Zhou et al.^45^ and (**) Yang et al.^46^.

### The transient expression of PsMYB10.2 promotes anthocyanin biosynthesis in the flesh of Japanese plum fruits

To determine whether the expression of *PsMYB10*.*2* promoted anthocyanin biosynthesis, its genomic sequence obtained from a red (H2/H2 genotype) and a yellow flesh (H4/H4) selections was transiently overexpressed in yellow flesh Japanese plum fruits. The fruits agroinfiltrated with either the *PsMYB10*.*2*-H2 or the *PsMYB10*.*2*-H4 alleles showed visible red stains on the yellow flesh in the region surrounding the injection (Figure 4), indicating that both coded proteins were functional. These red stains were not observed in other areas of the fruits, in fruits injected with the negative control vector (pBI121), and nor in noninjected fruits. The red stains in the flesh corresponded to a 20 to 103-fold increase of anthocyanin levels compared with the fruits injected with the control vector.

**Figure 4.**
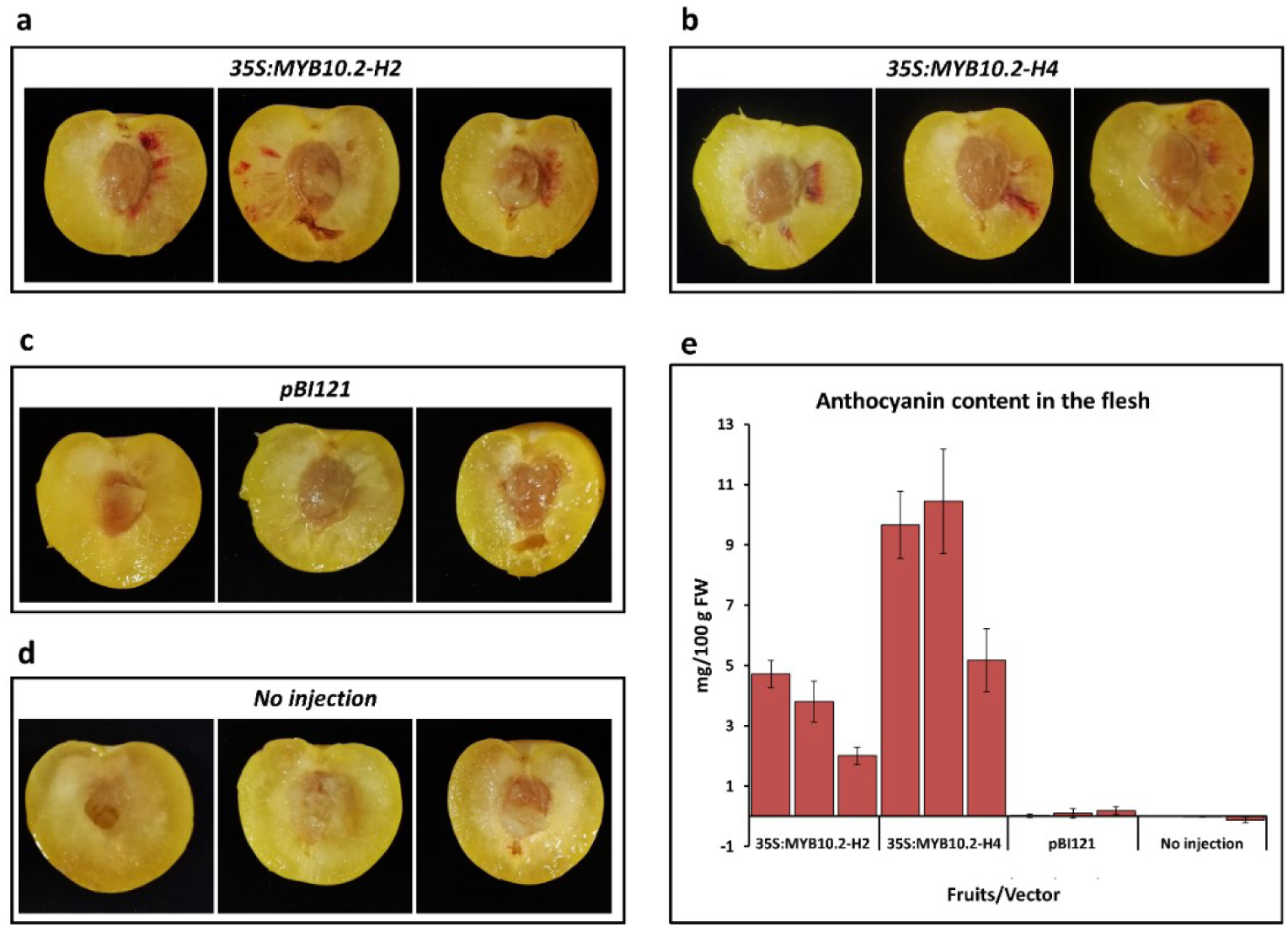
Transient overexpression of the *PsMYB10*.*2*-H2 and *PsMYB10*.*2*-H4 alleles in yellow flesh fruits and anthocyanin content. (**a**) fruits agroinfiltrated with the PsMYB10.2-H2 allele; (**b**) fruits agroinfiltrated with the PsMYB10.2-H4 allele. Negative controls: (**c**) fruits injected with the pBI121 vector and (**d**) noninjected fruits. (**e**) The anthocyanin content in the flesh of three fruits per treatment, error bars correspond to the standard deviation of each sample analyzed.

### An LTR retrotransposon is inserted in the PsMYB10.2-H2 promoter

To identify polymorphisms in the promoter that could be responsible for the lack of expression of the *PsMYB10*.*2* alleles in haplotypes other than H2, we did a BLAST search of the H1 and H2 allele sequences against the *de novo* assembly of the variety ‘Black Gold’ (H1/H2) obtained by Fiol et al. (submitted) through CRISPR-Cas9 enrichment sequencing. Only one contig for the *PsMYB10*.*2*-H2 allele was identified in ‘Black Gold’ (with 100% identity), with a sequence extending 3,142 bp upstream of the start codon. The BLAST search of this contig against all ‘Black Gold’ contigs identified an additional one 5,004 bp long. Both contigs overlapped 2,458 bp with 99.02% identity; the two sequences were merged to obtain a consensus sequence of 5,522 bp upstream of the *PsMYB10*.*2*-H2 gene start codon. For the *PsMYB10*.*2*-H1, we identified a contig with 99.7% identity in ‘Black gold’, which extended 9,339 bp upstream of the gene start codon. Alignment of the isolated H1 and H2 contigs revealed a 2.8 kb insertion in H2, 633 bp upstream of the gene start codon (Figure 5a). The BLAST search of this 2.8 kb sequence against the LG3-*PsMYB10* region assemblies of the five varieties from Fiol et al. (submitted) (‘Angeleno’ H1/H3, ‘Black Gold’ H1/H2, ‘Fortune’ H3/H6, ‘Golden Japan’ H4/H9, and ‘TC Sun’ H4/H5) gave positive hits in ‘Black Gold’ only. The BLAST search of the same sequence against the ‘Sanyueli’ and ‘Zhongli No.6’ reference genomes^35,48^ resulted in hits distributed along the eight chromosomes, none of them in the LG3-MYB10 region. Further search in the Rosaceae plants transposable elements database (RPTEdb) identified homology (E-value=0.0) with two *Prunus mume* Ty1-copia LTR retrotransposons: RLCopia_1_485_pmu and RLCopia_1_510_pmu. With the LTR_FINDER software, conserved sequences of long terminal repeat (LTR) retrotransposons were identified in the negative strand: the primer-binding site (PBS), the polypurine tract (PPT), the two LTR regions (762 bp and 772 bp long, with 97.4% of sequence identity), and the conserved 5’ TG and 3’ CA nucleotide pairs on them. The LTR retrotransposon insertion in the *PsMYB10*.*2* promoter was validated by long-range PCR. The amplification produced a band of 5 kb in a red flesh H2/H2 and of 2 kb in a yellow flesh H1/H1 selection (C31 and C27, respectively) (Figure 5b). The short band was also amplified in an H4/H4 selection (C12) and in C57, which was the only yellow flesh selection in the collection homozygous for the H2. Sequencing of the bands confirmed that the difference in size corresponded to the insertion of the LTR retrotransposon in H2 (Figure 5c). A characteristic of transposon insertions is the generation of a duplicated sequence at the genomic integration site, called Target Site Duplication (TSD). In the H2 sequence of the red flesh selection, a 5 bp TSD sequence was identified flanking the two LTR regions. This 5 bp sequence was found without duplication in the *PsMYB10*.*2* promoter of yellow flesh selections with H2 and H4, and was missing in the *PsMYB10*.*2*-H1 promoter.

**Figure 5.**
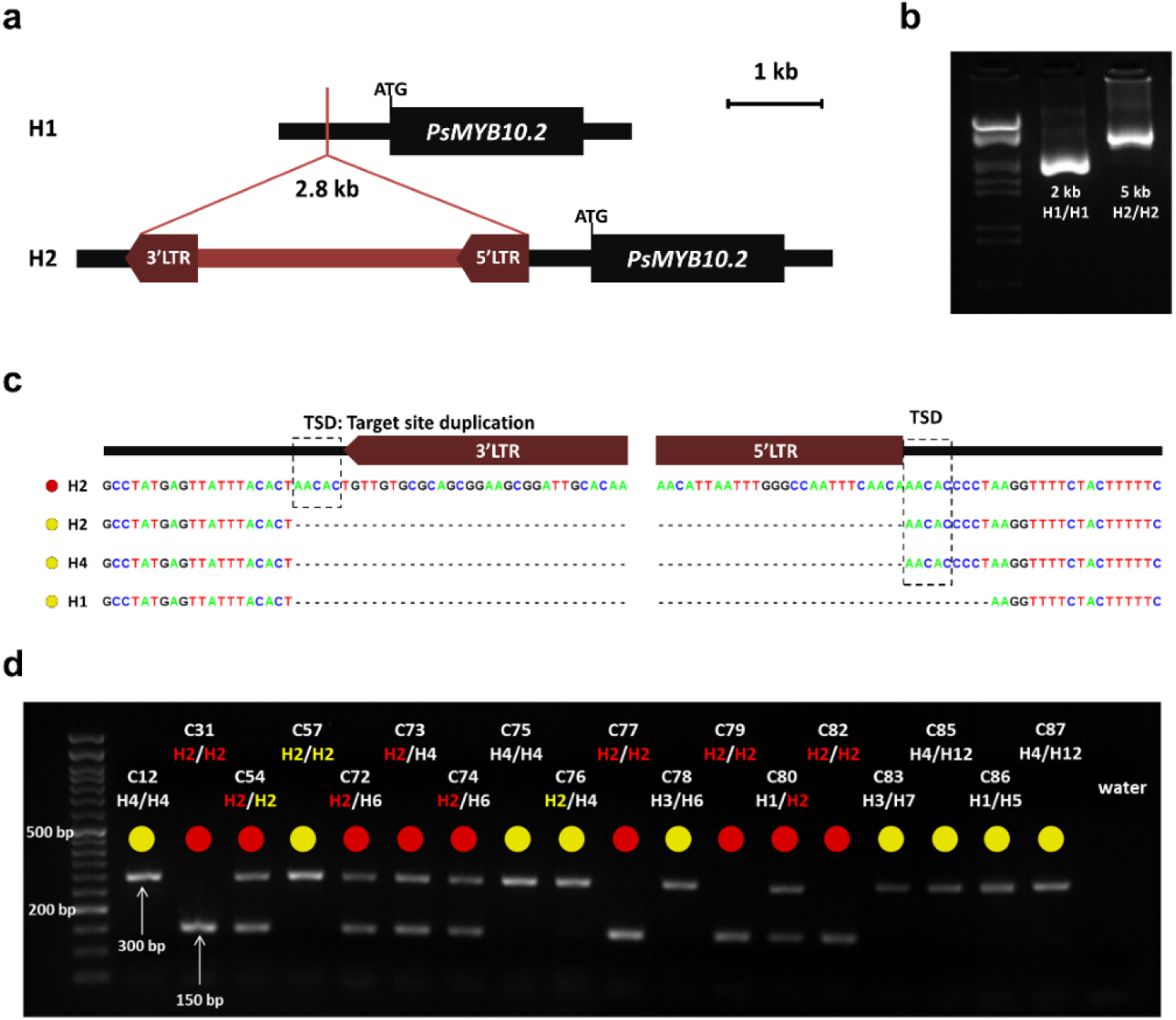
(**a**) Graphic representation of the 2.8 kb LTR retrotransposon insertion in the *PsMYB10*.*2* gene promoter in H2. (**b**) PCR amplification of the region in the *PsMYB10*.*2* promoter in C27 (H1/H1) and C31 (H2/H2) selections. The ladder in the first lane corresponds to Lambda DNA fully digested with HindIII and EcoRI enzymes. (**c**) Sequence of the retrotransposon insertion site in different haplotypes, the red and yellow circles correspond to red flesh and lack of red flesh, respectively.

The TSD (target site duplication) produced by the mechanism of the retrotransposon insertion is indicated. (**d**) Agarose gel visualization of a representative set of Japanese plum selections, the 150 bp band corresponds to the presence of the LTR retrotransposon and the 300 bp band to its absence.

### The presence of the LTR retrotransposon correlates with red flesh color

The insertion of the LTR retrotransposon in the *PsMYB10*.*2* promoter was evaluated in the collection of 103 Japanese plum selections with a set of PCR primers, giving a 150 bp band when the LTR retrotransposon was present and a 300 bp band when it was absent (Figure 5d; Supplementary Data 1). Among the 41 selections with red flesh, the LTR retrotransposon was found in either homozygosis or heterozygosis in all the 32 genotyped with H2 plus one H4/H8 individual (C43). The eight red selections without the LTR retrotransposon acquired the red color at the very late stage of ripening. These included selection 98-99 and five seedlings of the same parent (‘Sweet Pekeetah’, with yellow flesh). In the yellow flesh group, only C4 (H1/H2) had the LTR retrotransposon insertion. RT-PCR analysis showed that the *PsMYB10*.*2* gene was expressed in the mature flesh of C4, indicating that, in this selection, anthocyanin biosynthesis was repressed downstream in the pathway.

Our results indicate that there are two H2 haplotypes differing in the insertion of the LTR. While H2 selections lacking the LTR produce fruits without anthocyanins in their flesh, those with the LTR produce red-fleshed fruits. The LTR explained flesh color in 91.26% of the germplasm analyzed here. In all cases, the LTR insertion was linked to the *PsMY10*.*2* gene expression, suggesting a role of this polymorphism in the promotion of the gene expression. The LTR was also inserted in C43 (H4/H8) suggesting that one of the two haplotypes has also two forms: one with the LTR retrotransposon inserted, producing red fruits, and one without.

The association of the LTR retrotransposon with red flesh color was further validated in a collection of 42 commercial varieties (eight with red flesh and 34 with yellow or orange flesh). The LTR was able to predict the presence or lack of anthocyanin flesh color in 97.6% of them. The prediction only failed for ‘Black Beauty’ (yellow flesh, H1/H2), which, as C4, had the LTR insertion in heterozygosis (Supplementary Data 1). Considering all the individuals of the study, the retrotransposon explained the flesh color of 93.1% of the 145 genotypes analyzed.

## DISCUSSION

### Activation of the PsMYB10.2 gene promotes anthocyanin synthesis in the fruit flesh

In a previous study^23^ we isolated *PsMYB10* alleles and found one highly associated with the red skin color and none with the flesh color. A PCR primer combination was then developed for marker-assisted selection (MAS). The MAS method relied on the organization of the alleles into haplotypes. Here we increased the sample size and looked for correlation between the haplotypes instead of the alleles, with the red/non-red flesh trait, identifying a strong correlation of the H2 haplotype with the red flesh phenotype (p-value=8.06×10^−12^). This provides another example of the advantages of using haplotype data rather than only single polymorphisms^49-51^. However, the association did not explain the flesh color in 15.53% of the germplasm tested (consisting of advanced breeding lines and a few seedlings): these produced red flesh fruits without carrying the H2 haplotype or had the H2 haplotype but had yellow, orange or white flesh fruits. In Fiol et al.^23^ we found that one allele, a467, was always present in H6 and in a few individuals with H2. This allele was in the H2 of most of the yellow flesh individuals. We hypothesized here that H2 could have two versions: one associated with the red flesh color, and one (hereby defined as H2*) non-associated.

The expression of the *PsMYB10*.*2* allele in H2 individuals with red flesh suggested that this gene had a functional role in the synthesis of anthocyanins in the flesh. This is in agreement with Fang et al.^52^, who found that anthocyanin biosynthetic genes and the *PsMYB10*.*2* transcription factor were upregulated in a red flesh radiation-induced mutant of a yellow flesh Japanese plum variety. Here we show that the *MYB10*.*2* transcription factor is accountable for part of the natural variation of the trait. Fang et al.^52^ validated the gene function by transient overexpression in *Nicotiana*, co-infiltrating the MYB10.2 and the *PsbHLH3* genes (required to activate anthocyanin synthesis through the R2R3-MYB-bHLH-WD-repeat complex, MBW). We performed the transient overexpression in yellow Japanese plum fruits, which did not require co-expression of the *bHLH3* gene with the *PsMYB10*.*2* to observe the pigment accumulation in the flesh. This indicates that, in the samples tested, the mechanism required for activation of the anthocyanin biosynthetic genes was functional and that the main determinant for anthocyanin accumulation in the flesh was the expression of the *PsMYB10*.*2* gene.

Functional amino acid modifications and key residues have been described in the *MYB10*.*2* protein of peach^5,7,45,46^. In Japanese plum, there were no relevant differences of expressed and non-expressed alleles of the *PsMYB10*.*2* gene at the translated amino acid level, suggesting that the differences in their functionality could be due to polymorphisms in the promoter or in the gene non-coding regions. Consistent with this hypothesis, the overexpression of *PsMYB10*.*2-H2* and *PsMYB10*.*2-H4* alleles in yellow flesh fruits initiated the synthesis of anthocyanins. Since the *PsMYB10*.*2*-H4 allele was not expressed in fruits and its protein was the most distant to that of the PsMYB10.2-H2 allele, the results hinted that all the *PsMYB10*.*2* alleles cloned may be *per se* functional and that the observed differences in flesh coloration were due to the activation of the *PsMYB10*.*2-H2* gene.

### An LTR retrotransposon in the PsMYB10.2 allele promoter region may activate the gene expression

The CRISPR-Cas9 enrichment strategy applied in the *PsMYB10* region^44^ provided a reliable source of data to search for polymorphisms in the promoter of the *PsMYB10*.*2-H2* allele that could explain the gene expression. One of the advantages of long-read over short-read high-throughput sequencing technologies is the ability to detect structural variants^53^, which served here to locate a 2.8 kb retrotransposon inserted in the *PsMYB10*.*2-H2* gene promoter. This large polymorphism involving a highly repeated sequence in the genome might have been difficult, or even impossible, to identify by other means.

Transposable elements can modulate gene expression by introducing transcription factor binding sites, transcription initiation sequences or *cis*-regulatory motifs that respond to environmental factors. Together with other genetic and epigenetic mechanisms they have been found to be responsible for several cases of phenotypic variation in plants^54^. The retrotransposon identified here belongs to the long terminal repeat (LTR) family, which has already been reported to play a role in *MYB10* variability and the shift of anthocyanin fruit color in other rosaceous crops. The case reported in this study might be similar to that for red-skin apples, where an LTR retrotransposon in the *MdMYB10* promoter was associated and functionally validated with the increased gene expression^24^. In strawberry, an LTR retrotransposon disrupting the *MYB10* gene sequence was causative of the white-flesh phenotype, while in another allele a CACTA-like transposon inserted in the promoter was associated with the enhanced gene expression observed in the red fruits^14^. Outside the Rosaceae family, cases of transposons enhancing the expression of the orthologous R2R3-MYB gene have been reported in purple cauliflower, blood orange and pepper^55-57^.

In the advanced breeding line collection studied here, we observed that the LTR retrotransposon was dominant for the red flesh color and improved the initial correlation of H2 (the LTR explained the phenotype of 91% of the lines versus the 84% explained by the H2 haplotype). The number of alleles shared between H2 and H2* and the TSD identified in H2 suggest that the latter originated after the insertion of the LTR retrotransposon in the original *PsMYB10*.*2*-H2* allele. Further functional studies are required to verify if the retrotransposon is the main determinant enhancing *PsMYB10*.*2*-H2 gene expression and to locate regulatory sequences within the transposable element or within the disrupted insertion site.

### PsMYB10 genes make a great contribution to the Japanese plum skin and flesh colors

As for the advanced breeding lines, the fruit skin and flesh color in the panel of 42 commercial varieties were largely explained by the variability in the LG3-*MYB10* genes (only one outlier for the flesh color, none for the skin color markers), which confirmed the strong association of the region with the fruit color trait and the usefulness of the molecular markers designed. The MYB10 region contains several gene copies in tandem^44^, a conformation that could have been generated by unequal crossing over by either homologous or non-homologous recombination^58^.

After a gene duplication event, the subsequent fate of the copy can be i) conservation of the gene function (maintenance), ii) degeneration into a pseudogene or its removal (non-functionalization), iii) very rarely, the gain of a novel function (neo-functionalization) or, together with the original gene, iv) the accumulation of mutations which subdivide their functionality, complementing each other to achieve the function of the ancestral gene (sub-functionalization)^59,60^. This sub-functionalization can be generated by changes in the gene regulatory sequences, altering their spatial and temporal expression^58^. This is likely the case in the *PsMYB10*.*1* and *PsMYB10*.*2* genes which greatly differ in their upstream regulatory sequences, causing one to be expressed in the skin and the other in the flesh, respectively, localizing anthocyanin synthesis in different fruit tissues. The haplotypes observed by Fiol et al.^23^ (based on the segregation of the alleles of the *PsMYB10*.*1, PsMYB10*.*2* and *PsMYB10*.*3* gene copies) dominant for the red skin (H1, H3; both with the *PsMYB10*.*1*-356 allele) are recessive for yellow flesh, while the haplotype dominant for the red flesh color (H2; with an LTR in the promoter of the *PsMYB10*.*2* gene) is recessive for the yellow skin (Figure 6). As such, the presence of H2 with either H1 or H3 is required to confer anthocyanin coloration in both tissues. Okie (2008)^61^ described that plums with yellow skin were transparent and if accompanied by red flesh they were mottled and bronze-like, matching the fruits in our study with background skin color (or mottled-type) and red flesh, which have H2 and any except H1 or H3.

**Figure 6.**
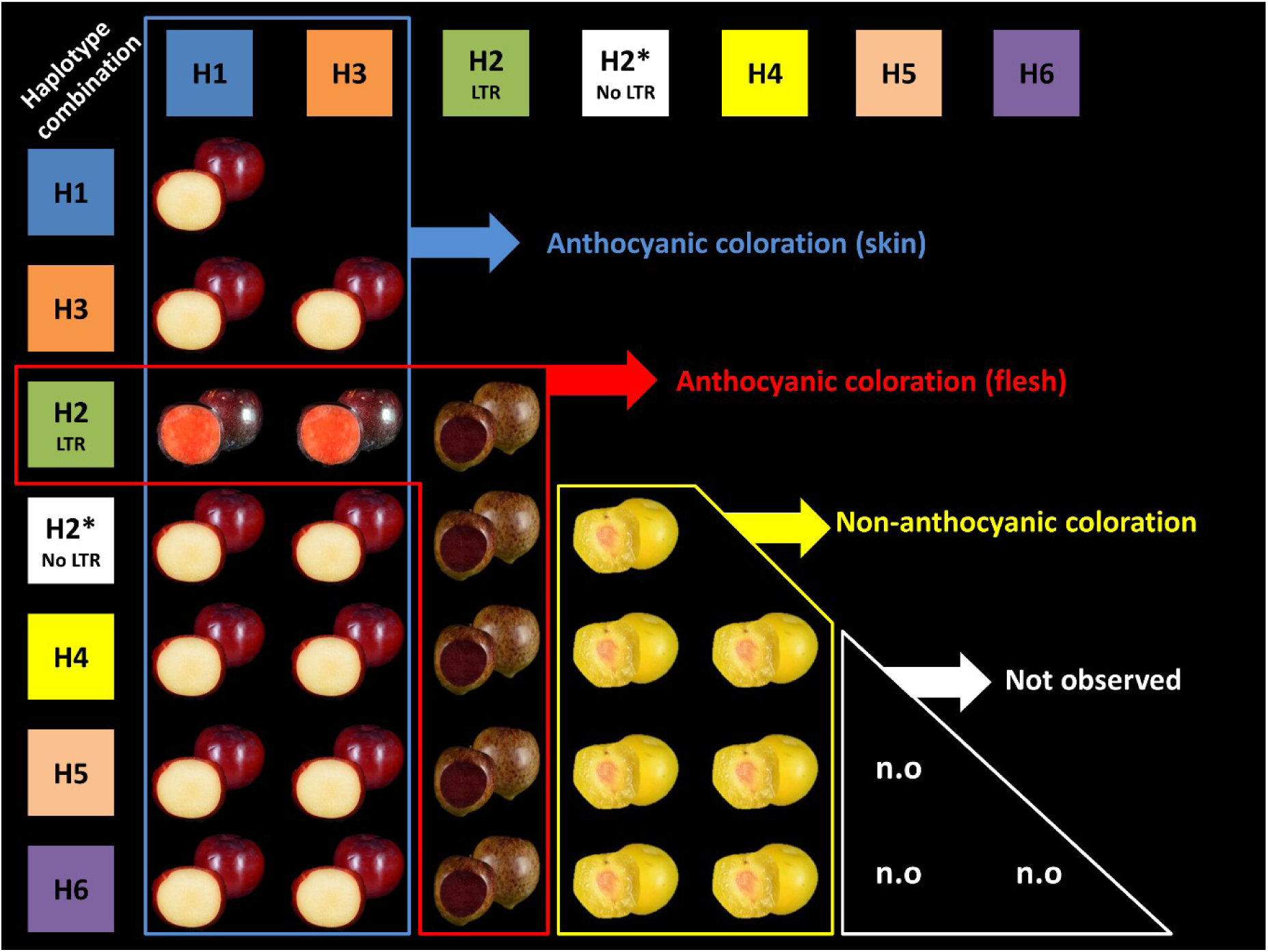
Japanese plum fruits showing the presence or absence of anthocyanin color in skin and flesh for every possible combination of the seven observed and most frequent MYB10 haplotypes found in the collections of Japanese plum selections and commercial varieties. The combinations H1/H2 and H2/H3 produce anthocyanins in the skin and flesh. We did not observe combinations with H5 and H6, but we expect they lack anthocyanins in both tissues.

### Not all the fruit flesh color variability in Japanese plum is attributable to the identified MYB10 haplotypes

Fiol et al.^44^ studied the synteny between the LG3-MYB10 region in the genome assemblies of *P. salicina* (two genomes), *P. avium* (two genomes), *P. persica* (two genomes), *P. mira, P. armeniaca* (five genomes), *P. sibirica* (two genomes) and *P. mandshurica*, demonstrating the high complexity of this region in Japanese plum, with homology levels between varieties comparable to those between *Prunus* species. This, together with the identification here of two H2 haplotypes (H2 and H2*) plus the insertion of the LTR in either H4 or H8 in C43 suggest that there may be additional polymorphisms in the LG3-*PsMYB10* region that could account for the variability in the flesh color not explained by the LTR.

In addition, similarly to what has been reported in peach, a single genomic region may not be responsible for all the red flesh color variability in Japanese plum. In the 103 selections, the only one non-anthocyanic with the LTR was C4 (H1/H2). When this line was crossed with another yellow selection (C14 [H3/H4]), some offspring (all with H2) had red flesh while, as with C4, some offspring with H2 were yellow^23^. We observed a similar segregation in the first year of fruiting of F1 progeny of C4 with C11 (yellow flesh, H3/H6) (data not shown), indicating that C4 can be a donor of the red flesh color trait through the LTR retrotransposon despite being yellow. Interestingly, the *PsMYB10*.*2* gene is expressed in the C4 flesh, which could be explained by the presence of a downstream regulator of the anthocyanin biosynthesis pathway. The color prediction in the commercial cultivar collection failed only in ‘Black Beauty’, also H1/H2, which could be a similar case.

A detailed observation of the trait showed that the phenotype in most of the red individuals without the LTR insertion was different to that observed in those with the insertion. While selections and varieties such as ‘Black Gold’ and ‘Black Splendor’ (H1/H2) accumulated the anthocyanin color progressively and homogeneously in the flesh during ripening (Supplementary Figure 1a), others without the insertion acquired color in the very late stages of fruit ripening and firstly around the skin. One example is selection 98-99 considered here as red flesh as in^42^ although has been also described as yellow flesh^31,62,63^ (Supplementary Figure 1b). Other five red-fleshed individuals without the LTR are offspring of ‘Sweet Pekeetah’, which does not have the retrotransposon and has yellow flesh: the late red flesh color might segregate through other genomic region/s. We noted that, in some cases, the red pigmentation of the flesh was only localized near the red or black skin (Supplementary Figure 1c). While this could be due to the direct effect of another genomic region from that studied here, a possible contribution of the skin color to that of the flesh should not be discarded. Also not explained by the LTR retrotransposon insertion is the synthesis of anthocyanins induced by chilling stress (5°C). This phenotype has been reported in some yellow flesh commercial cultivars studied here: ‘Friar’ (H1/H5), ‘Black Amber’ (H1/H3) and ‘Royal Diamond’ (H1/H3)^64-66^.

Additional QTL studies are required to identify more genomic regions involved in the regulation of flesh coloration. Due to appearing earlier in the maturation process, the homogenous red flesh color trait, if expressed, could have an epistatic effect over the presence of late coloration. As also noted by other authors, it will be interesting to study not only the presence of red flesh color but when^42^ and where it appears, which would facilitate identifying other genomic regions controlling fruit coloration localized in specific fruit tissues and appearing in different ripening stages and/or environmental conditions.

## CONCLUSION

In this study we found that most of the natural variability observed in the Japanese plum fruit flesh color is triggered by the expression of the *PsMYB10*.*2* gene. We have shown that when the *PsMYB10*.*2* gene is expressed in the flesh of mature fruits, the biosynthesis of anthocyanin is activated and the red pigment accumulates in the flesh. As has been demonstrated in other rosaceous species, the insertion of an LTR retrotransposon in the promoter of the gene may promote its expression. Apart from contributing to the knowledge of the genetic mechanisms behind the natural variation of Japanese plum fruit color, here we provide i) an improved protocol for the transient gene overexpression mediated by *Agrobacterium* in the flesh of Japanese plum fruits, that can be used for the validation of other candidate genes; ii) an efficient molecular marker for the flesh color which, together with the one previously developed by our group for skin color, can be used for marker-assisted breeding. However, as previously discussed the anthocyanin synthesis is complex, regulated by a network involving additional loci and environmental factors, and all the flesh color variability may not be determined by a single region. Therefore, further studies are needed to capture phenotypic variation caused by other loci.

## MATERIALS AND METHODS

### Plant material and nucleic acid isolation

A collection of 103 Japanese plum selections from two breeding programs was analyzed, consisting of 81 trees cultivated in Huelva (Spain) and 22 trees cultivated in Rinconada de Maipú (Chile) (Supplementary Data 1). Fruit color was visually phenotyped from several fruits at maturity for a minimum of two seasons. Flesh color was categorized as white, green, yellow, orange or red; skin color was classified as mottled, green, yellow, red, purple or black.

The fruits from five cultivars with different flesh color were obtained in triplicate at maturity from the local market. These were ‘Angeleno’ (pale yellow flesh), ‘Golden Japan’ (yellow flesh), ‘Rose’ (orange flesh), ‘Black Gold’ (red flesh) and a mottled-type plum (dark red flesh). The skin was removed and the flesh tissue was used for RNA and DNA extraction. The mature flesh of three fruits from the selection C4 was also used for RNA extraction. Transient gene expression assays were carried out using 30 yellow flesh fruits of ‘Golden Japan’ cultivar, harvested one week before full maturity stage.

To further validate the results, a DNA collection of 42 commercial varieties was analyzed (Supplementary Data 1). The DNA was extracted from the leaves in all cases except for ‘Rose’ and the mottled-type plum, where the DNA was extracted from the fruit. Fruit color descriptors for this collection were obtained from breeders’ descriptions.

DNA was extracted following the CTAB procedure^67^. The RNA was extracted with the Maxwell RSC Plant RNA Kit (Promega) and was further DNase-treated with the TURBO DNA-free Kit (Invitrogen). The quality of the extracted nucleic acids was assessed using a Nanodrop ND-1000 Spectrophotometer and 0.8% agarose gels. All the studied material, including their fruit color phenotype and their genotyping results, can be found in Supplementary Data 1.

### MYB10 genotyping and association with flesh color

All extracted DNAs were analyzed with the MYB10 markers (MYB10F2/MYB10NR2 and MYB10F2/MYB10NR4 primer pairs) to obtain their haplotype combination^23,68^. A χ^2^ test was run on the collection of 103 Japanese plum selections to correlate the H1 to H6 haplotypes with the presence or absence of anthocyanin-based coloration of the fruit flesh. For this, it was assumed that the red phenotype category indicated anthocyanin presence and the remaining categories (white, green, yellow and orange) its absence.

### Expression of the MYB10 genes in the fruit flesh

The RNA samples extracted from the fruit flesh were reverse transcribed using the PrimeScript RT-PCR Kit (TaKaRa) and the oligo(dt)20 primer. The expression of the MON reference gene^69^ was used for quality control of the synthesized cDNAs. *PsMYB10*.*1, PsMYB10*.*2* and *PsMYB10*.*3* gene expression in the flesh of the red ‘Black Gold’ and the yellow ‘Golden Japan’ cultivars was evaluated by PCR. Primers and annealing temperatures used for each gene are described in Supplementary Table 2. The PCR reactions were carried out using 1.5 mM MgCl_2_ and 1X NH_4_ buffers for 1U of BioTaq polymerase (Bioline), 0.2 mM dNTP mix, 0.2 µM of each primer, 2 µl of cDNA and MilliQ water adjusted to a 10 µl reaction. PCR conditions were 1 minute at 94°C, 35 cycles of 94°C for 30 seconds, the annealing temperature of each primer pair for 20 seconds and 72°C for 2 minutes, followed by a final elongation step of 5 minutes at 72°C. Genomic DNA extracted from the fruit of each variety was added as amplification control. Amplification results were visualized in 2% agarose TAE gels after running 45 minutes at 120V. Amplification from the cDNAs only occurred with ‘Black Gold’ *PsMYB10*.*2*. The same reaction was repeated adding the cDNAs of ‘Angeleno’, ‘Rose’ and the mottled-type plums, and later repeated with the cDNA of C4. The *PsMYB10*.*2* gene product from ‘Black Gold’ was purified using PureIT ExoZAP Kit (Ampliqon) and sequenced by capillary electrophoresis with the same primers used to amplify the fragment.

### PsMYB10.2 sequencing on different haplotypes

The *PsMYB10*.*2* alleles on haplotypes H1 to H4 were PCR-amplified from the genomic DNA of *PsMYB10* homozygous selections using the methodology described above. The selections were C3 (H1/H1), C31 (H2/H2), C6 (H3/H3) and C12 (H4/H4), being C31 the unique with red flesh color. The PCR purified products were ligated to the pCR™2.1-TOPO® vector (Invitrogen), following the manufacturer’s instructions, and sequenced. The obtained sequences were aligned to the *Prunus salicina* MYB10 gene from the NCBI database (EU155161.1) and the *PsMYB10*.*2* cDNA sequence from ‘Black Gold’ using Sequencher 5.0 software (Gene Codes Corporation, Ann Arbor, MI, USA). The introns were removed, and the *in-silico* coded protein sequences obtained with the ExPASy Translate tool^70^ were aligned to the peach *PpMYB10*.*2* (*Prupe*.*3G163000*) amino acid sequence using Clustal Omega^71^ and visualized using Jalview v2^72^.

### Vector construction

The genomic 2 kb sequences from *PsMYB10*.*2*-H2 and *PsMYB10*.*2*-H4 were PCR-amplified from the C31 and C12 selections, respectively, using the Phusion Green High-Fidelity DNA Polymerase kit (Thermo Scientific) with primers that included restriction enzyme sites (Supplementary Table 2). The reactions were purified with the High Pure PCR product purification kit (Roche) and digested with SacI and XbaI enzymes in M buffer (TaKaRa). The gene sequence included a SacI restriction site in the second intron and, after 1 hour of XbaI digestion, the amplicon was partially digested with SacI by limiting the reaction to 2 minutes at 37°C. The pBI121 vector was fully digested with the same enzymes to remove the GUS gene, then the genic sequences were ligated next to the CaMV 35S promoter using T4 DNA Ligase (Thermo Scientific) to construct the 35S:MYB10.2-H2 and 35S:MYB10.2-H4 vectors. The ligation products were transformed into *Escherichia coli* JM109 competent cells and grown in LB media supplemented with 50 mg/l of kanamycin; screening was performed by direct colony PCR using M102_ex3_F and M13fwd primers (Supplementary Table 2). The vectors from positive colonies were extracted with the GenJet Plasmid Miniprep kit (Thermo Scientific) and sequenced before transformation into *Agrobacterium tumefaciens* EHA105 competent cells.

### Transient gene expression in Japanese plum fruits

The *MYB10*.*2* function was validated by its transient overexpression in ripe fruits. The procedure was based on previous studies describing the genetic transformation in plum and the transient expression in fruits of closely related species^73-76^, with some modifications to increase its efficiency. Specifically, *Agrobacterium tumefaciens* EHA105 cells transformed with the pBI121, the 35S:MYB10.2-H2 and the 35S:MYB10.2-H4 vectors were grown for 24 hours in liquid LB media with kanamycin and rifampicin (50 mg/l), then 1 ml of each culture was used to inoculate 15 ml of LB liquid media containing 20 µM of acetosyringone and 10 mM MES (2-(N-Morpholino)ethanesulfonic acid) at pH 5.6, and grown overnight. The final OD_600_ of the culture was quantified in a UV-2600 spectrophotometer (Shimadzu), then cells were pelleted by centrifugation and resuspended to a OD_600_ of 0.8 in injection buffer (10 mM magnesium chloride hexahydrate, 200 µM acetosyringone, 10 mM MES at pH 5.6) and kept in the dark at room temperature for 2-4 hours with occasional shaking. The 30 ‘Golden Japan’ fruits used for the experiment were rinsed 2 minutes in a 30% bleach solution, washed with Tween-20 (0.05%) and completely dried. Ten fruits were injected with the *Agrobacterium* solution carrying the 35S:MYB10.2-H2 vector, ten more with the 35S:MYB10.2-H4, five with the pBI121, and the remaining five fruits were not injected. Using 1 ml hypodermic syringes, a dose of 1 ml was evenly distributed on one side of each fruit by following the suture line, with a minimum of three injection points per fruit. Fruits were kept at room temperature in the dark and opened in halves after 13 days by cutting from the suture line. Anthocyanin was quantified using the differential pH spectrophotometry method as used in Fang et al.^77^ in Japanese plum fruits, weighing approximately 0.1 g of flesh tissue extracted from the observable red flesh patches. The same approximate quantity was extracted from the injection sites in the case of the pBI121-treated fruits and randomly in the noninjected group of fruits, which served as negative controls.

### Identification of polymorphisms in the PsMYB10.2-H2 promoter

To identify polymorphisms in the *PsMYB10*.*2* promoter, the CRISPR-Cas9 MYB10-enriched MinION sequencing data from Fiol et al.^44^ was investigated. This data included sequences from the red flesh cultivar ‘Black Gold’ (H1/H2) and from the non-red flesh varieties ‘Angeleno’ (H1/H3), ‘Fortune’ (H3/H6), ‘Golden Japan’ (H4/H9) and ‘TC Sun’ (H4/H5). The blastn software^78^ was run with the H1 and H2 *PsMYB10*.*2* input sequences in the ‘Black Gold’ assembly, selecting the contigs with the best score hits that extended upstream of the gene. The homologous *PsMYB10*.*2*-H2 contig ended prematurely and was extended by selecting the only contig that overlapped the upstream sequence after a blast analysis. By aligning and comparing the sequences in Sequencher 5.0, a large insertion on the H2 contig compared to the H1 was identified. This sequence was blast searched in the CRISPR-MinION sequences of the five varieties, and then on the *P. salicina* ‘Sanyueli’ v2.0^48^ and *P. salicina* ‘Zhongli No 6’ v1.0^35^ genomes using blastn software from GDR. The Blast search was repeated in the Rosaceae transposable elements database from the RPTEdb^79^ and the sequence was further analyzed with LTR_FINDER software^80^ to identify conserved structures from the LTR-type retrotransposon identified.

### Validation of the insertion with long range PCR and marker design

To validate the large insertion, primers pM102F and pM102R (Supplementary Table 2) were designed and used in a long-range PCR reaction using the LongAmp® Taq DNA Polymerase (NEB) to amplify 40 ng of genomic DNA following the kit instructions. The red flesh selection used was C31 (H2/H2), the yellow flesh selections were C27 (H1/H1), C57 (H2/H2) and C12 (H4/H4). The PCR amplicons were run in a 2% agarose gel for 45 minutes at 120V. The PCR products were purified and sequenced by capillary electrophoresis as described above. The sequencing results were aligned with Sequencher 5.0 to validate the insertion site and used in Primer3 software^81^ to design two primers flanking the 2.8 kb insertion on each side (LTR_F, LTR_R) and one within it (LTR_i) (Supplementary Table 2). The PCR reaction for the LTR marker was carried out using 0.2 µM, 0.4 µM and 0.15 µM of LTR_F, LTR_i and LTR_R primers, respectively, mixed with 1.5 mM MgCl_2_ and 1X NH_4_ buffers, 1U of BioTaq polymerase, 0.2 mM dNTPs and MilliQ water to 10 µl. Thermocycler conditions were 94°C for 1 minute, 35 cycles of 94°C for 15 seconds, 56°C for 30 seconds and 72°C for 15 seconds, followed by one step at 72°C for 5 minutes. This reaction was used to genotype for the absence or presence of the retrotransposon in all the extracted DNA material, loading the 10 µl PCR results into TAE gels with 2% agarose which run for 40 minutes at 120V.

## ACKNOWLEDGMENTS

AF is recipient of grant BES-2016-079060 funded by MCIN/AEI/10.13039/501100011033 and by “ESF Investing in your future”. CD was supported by “DON CARLOS ANTONIO LOPEZ” Abroad Postgraduate Scholarship Program, BECAL-Paraguay. This research was supported by project RTI2018-100795-B-I00 funded by MCIN/AEI/10.13039/501100011033 and by “ERDF A way of making Europe”. We acknowledge support from the CERCA Programme (“Generalitat de Catalunya”), and the “Severo Ochoa Programme for Centres of Excellence in R&D” 2016-2019 (SEV-2015-0533) and 2020-2023 (CEX2019-000902-S) both funded by MCIN/AEI /10.13039/501100011033.

We thank PLANASA company and their breeders Mario Ortiz and Antonio García for providing plant material and phenotype information. We also wish to thank Jorge Naranjo (Tany Nature), Alfonso Guevara (IMIDA) and David Ruiz (CEBAS-CSIC) for providing plant material from commercial varieties, and L. Maria Lois for providing the pBI121 vector.

## Notes

### Competing Interest Statement

The authors have declared no competing interest.

