## Supplementary figures and images for "An LTR retrotransposon in the promoter of a *PsMYB10.2* gene associated with the regulation of fruit flesh color in Japanese plum"

### Supplementary Figure 1

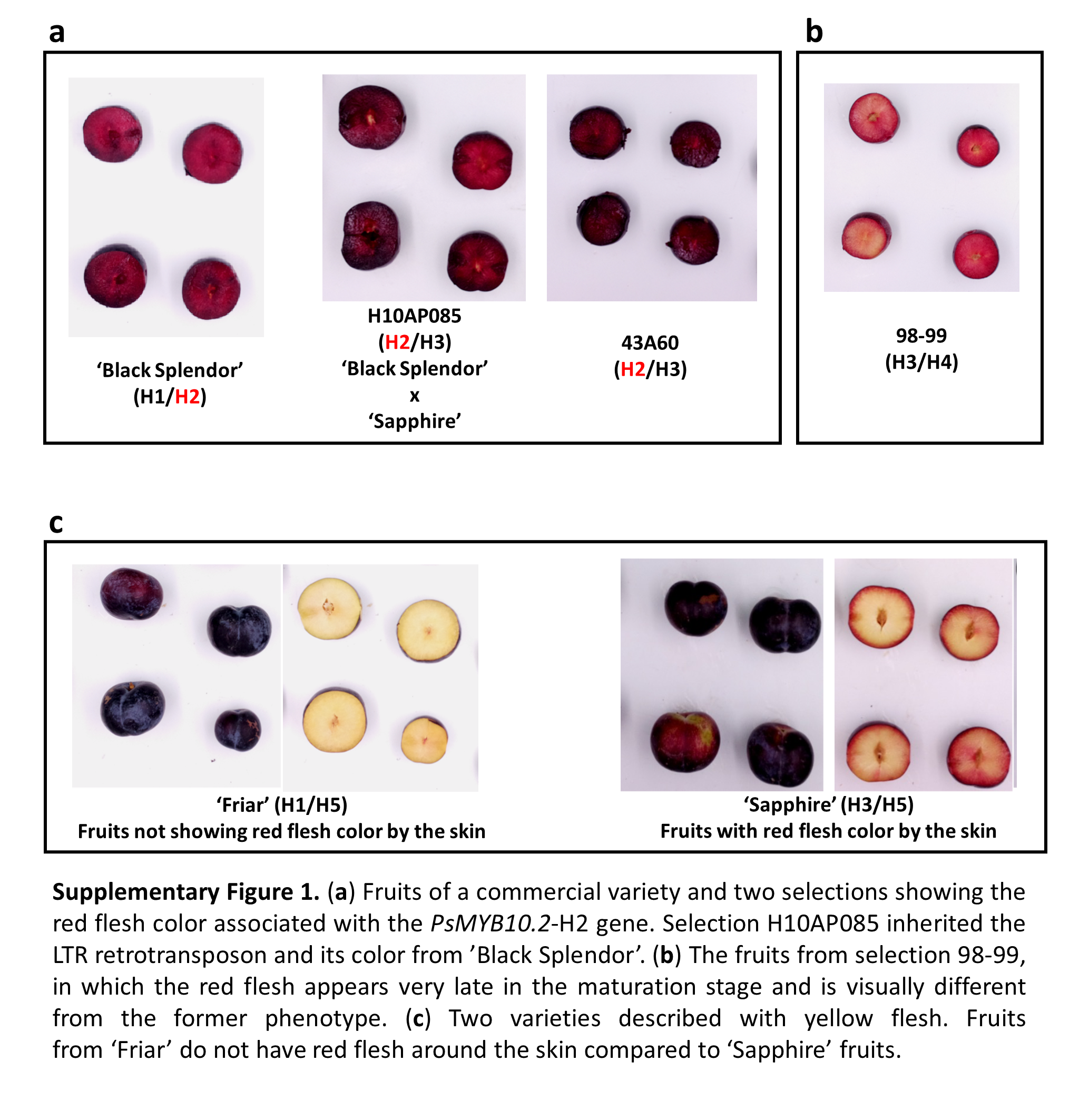
