## Supplementary Table 1 for "An LTR retrotransposon in the promoter of a *PsMYB10.2* gene associated with the regulation of fruit flesh color in Japanese plum"

| Haplotype | Frequency (%) | Chi-square p-value |
| --- | --- | --- |
| H1 | 21.36 | 9.58×10^-1^ |
| H2 | 21.84 | 8.06×10^-12^ |
| H3 | 17.48 | 1.76×10^-1^ |
| H4 | 17.96 | 1.31×10^-2^ |
| H5 | 3.88 | 3.73×10^-1^ |
| H6 | 6.80 | 7.37×10^-1^ |

**Supplementary Table 1.** The results for the χ^2^_(1df)_ test in the panel of 103 selections of the breeding lines collection, considering the six most frequent haplotypes (H1 to H6) and the flesh color. The presence of H2 was statistically associated with the presence of red color.
