## Supplementary Table 2 for "An LTR retrotransposon in the promoter of a *PsMYB10.2* gene associated with the regulation of fruit flesh color in Japanese plum"

**Supplementary Table 2.** Primer sequences and their temperatures of annealing (Ta) used in the PCR reactions.

| **Primer name** | **Sequence** | **Ta (ºC)** | **Description** |
| --- | --- | --- | --- |
| **MYB10F2** | GTGTGAGAAAAGGAGCTT | 55 | *MYB10* marker (Fiol et al., 2021) |
| **MYB10NR2** | GATATTTGGCTTCAAATAGTTC |  |  |
| **MYB10NR4** | TTCCTGCACCTGTTCAAC |  |  |
| **M101_RT_F** | TGGACACGAGACATTGCACG | 57 | *PsMYB10.1* amplification (Fiol et al., 2021) |
| **M101_RT_R** | CAATGGTCTTTTGACAGCCGC |  |  |
| **M102_f** | CTGGCTGCAAGCATAC | 57 | *PsMYB10.2* amplification (Fiol et al., 2021) |
| **M102_r** | GTGGGACAAACACTCTC |  |  |
| **M103_f** | ATAGGAACTAGCAGGCAC | 57 | *PsMYB10.3* amplification (Fiol et al., 2021) |
| **M103_r** | AGTTGCTAATAATTGCTACTAGG |  |  |
| **teM102_F2** | ACGCATCTAGAACGCTGCACAAAAGAAAACA | 62 | Primers including XbaI and SacI restriction sites for *PsMYB10.2* amplification and insertion into pBI121 vector |
| **teM102_R2** | TCAGTGAGCTCGTCCCTTAACTTTTCAATGTGGG |  |  |
| **M102_ex3_F** | ACAGGTTCTGGTCTTGGGTT | 55 | Screening of positive colonies carrying the *PsMYB10.2* insert |
| **M13fwd** | GTAAAACGACGGCCAGT |  |  |
| **pM102F** | TGTTAGGCTGAAATGCAGGA | 56 | Amplification of the *PsMYB10.2* promoter using a long-range PCR |
| **pM102R** | GTATGCTTGCAGCCAG |  |  |
| **LTR_F** | (*)TGAGTTAGGTTGCCTATGAGT | 57 | Molecular marker of the LTR retrotransposon. (*)can be fluorescently labeled for capillary electrophoresis |
| **LTR_i** | CGGCGCAGTATAAATCCTTG |  |  |
| **LTR_R** | TTTTCGTTGATTGTTTGTCCAA |  |  |
